# Identification of a Nervous System Gene Expression Signature in Colon Cancer Stem Cells Reveals a Role for Neural Crest Regulators *EGR2* and *SOX2* in Tumorigenesis

**DOI:** 10.1101/2021.02.02.428317

**Authors:** Joseph L. Regan, Dirk Schumacher, Stephanie Staudte, Andreas Steffen, Ralf Lesche, Joern Toedling, Thibaud Jourdan, Johannes Haybaeck, Nicole Golob-Schwarzl, Dominik Mumberg, David Henderson, Balázs Győrffy, Christian R.A. Regenbrecht, Ulrich Keilholz, Reinhold Schäfer, Martin Lange

## Abstract

Recent data support a hierarchical model of colon cancer driven by a population of cancer stem cells (CSCs). Greater understanding of the mechanisms that regulate CSCs may therefore lead to more effective treatments. Serial limiting dilution xenotransplantation assays of colon cancer patient-derived tumors demonstrated ALDH^Positive^ cells to be enriched for tumorigenic self-renewing CSCs. In order to identify CSC modulators, we performed RNA-sequencing analysis of ALDH^Positive^ CSCs from a panel of colon cancer patient-derived organoids (PDOs) and xenografts (PDXs). These studies demonstrated CSCs to be enriched for embryonic and neural development gene sets. Functional analyses of genes differentially expressed in both ALDH^Positive^ PDO and PDX CSCs demonstrated the neural crest stem cell (NCSC) regulator and wound response gene *EGR2* to be required for CSC tumorigenicity and to control expression of homeobox superfamily embryonic master transcriptional regulator *HOX* genes and the embryonic and neural stem cell regulator *SOX2*. In addition, we identify *EGR2, HOXA2, HOXA4, HOXA5, HOXA7, HOXB2, HOXB3* and the tumor suppressor *ATOH1* as new prognostic biomarkers in colorectal cancer.

## INTRODUCTION

Colorectal cancer (CRC), the third most common cancer and fourth most common cause of cancer deaths worldwide^1^, is a heterogeneous tumor driven by a subpopulation of CSCs, that may also be the source of relapse following treatment^2–5^. Elucidation of the mechanisms that regulate CSC survival and tumorgenicity may therefore lead to novel treatments and improved patient outcomes.

CSCs are undifferentiated cancer cells that share many of the attributes of stem cells, such as multipotency, self-renewal and the ability to produce daughter cells that differentiate^2,6,7^. Stem cells are controlled by core gene networks that include the embryonic master transcriptional regulator *HOX* genes^8,9^ and *SOX2*^10^, whose misregulation can result in aberrant stem cell function, developmental defects and cancer^11,12^. These genes are crucial for embryonic development and their expression is maintained in adult tissue stem cells, where they regulate self-renewal and differentiation^9,13–15^. *HOX* genes and *SOX2* are aberrantly expressed in several cancers, including CRC, and emerging evidence demonstrates their involvement in the transformation of tissue stem cells into CSCs^11,16–23^. Modulation of *HOX* genes and *SOX2* could therefore provide novel therapeutic strategies to block tumorigenesis and overcome therapy resistance in CRC and other CSC driven cancers.

During embryonic development of the neural crest, which gives rise to the peripheral nervous system (PNS) and several non-neuronal cell types^24^, *HOX* and *SOX* genes are regulated by retinoic acid^25,26^, a product of the normal tissue stem cell and CSC marker aldehyde dehydrogenase (*ALDH1A1, ALDH1A2, ALDH1A3*)^8,26–29^, and by the neural crest stem cell (NCSC) zinc finger transcription factor and wound response gene *EGR2* (*KROX20*)^30–41^.

Here we carried out whole transcriptome analysis of functionally tested ALDH^Positive^ CSCs from a panel of colon PDOs and PDX models and show that colon CSCs and Lgr5^Positive^ intestinal stem cells (ISCs) are highly enriched for nervous system development and neural crest genes. Furthermore, we demonstrate that the neural crest stem cell (NCSC) gene *EGR*2 is a marker of poor prognosis in CRC and modulates expression of *HOX* genes and *SOX2* in CSCs to regulate tumorigenicity and differentiation.

## RESULTS

### Colon cancer PDOs are heterogeneous and enriched for ALDH^Positive^ self-renewing CSCs

Colon cancer PDO models were established from freshly isolated primary tumors and metastases from colon cancer patients (Table S1) by embedding in growth-factor reduced Matrigel and cultivating in serum free media, as previously described^42–44^. Immunostaining of PDOs for the structural proteins EZRIN and EPCAM demonstrated that PDOs retain the apical-basal polarity and structural adhesion of the normal intestine (Figure 1A). Immunostaining of PDOs and equivalent PDX models for stem cell regulator Wnt signaling protein BETA-CATENIN demonstrated differences in nuclear localization of BETA-CATENIN and confirmed previous data demonstrating heterogeneous Wnt signaling activity within the tumors^43^ (Figure 1B). Increased aldehyde dehydrogenase (ALDH) activity, as measured using the Aldefluor™ assay, is a marker of CSCs in colon cancer and many other cancer types^29^. We previously carried out limiting dilution serial xenotransplantation of ALDH^Negative^ and ALDH^Positive^ cells and demonstrated that colon CSCs are ALDH^Positive^ and enriched for Wnt signaling activity^43^. However, ALDH^Negative^ cells also gave rise to tumors when transplanted at higher cell numbers. In order to determine if ALDH^Negative^ and ALDH^Positive^ cells maintained their self-renewal and tumorigenic capacity, we performed additional rounds of limiting dilution serial xenotransplantation of ALDH^Negative^ and ALDH^Positive^ cells (Figure 1E). These data confirmed that PDOs are enriched for ALDH^Positive^ cells compared to equivalent PDX models (Figure 1C and D) and that ALDH^Positive^ CSCs self-renew to maintain their tumorigenic capacity over extended rounds of xenotransplantation, but that ALDH^Negative^ cells do not (Figure 1E).

**Figure 1.**
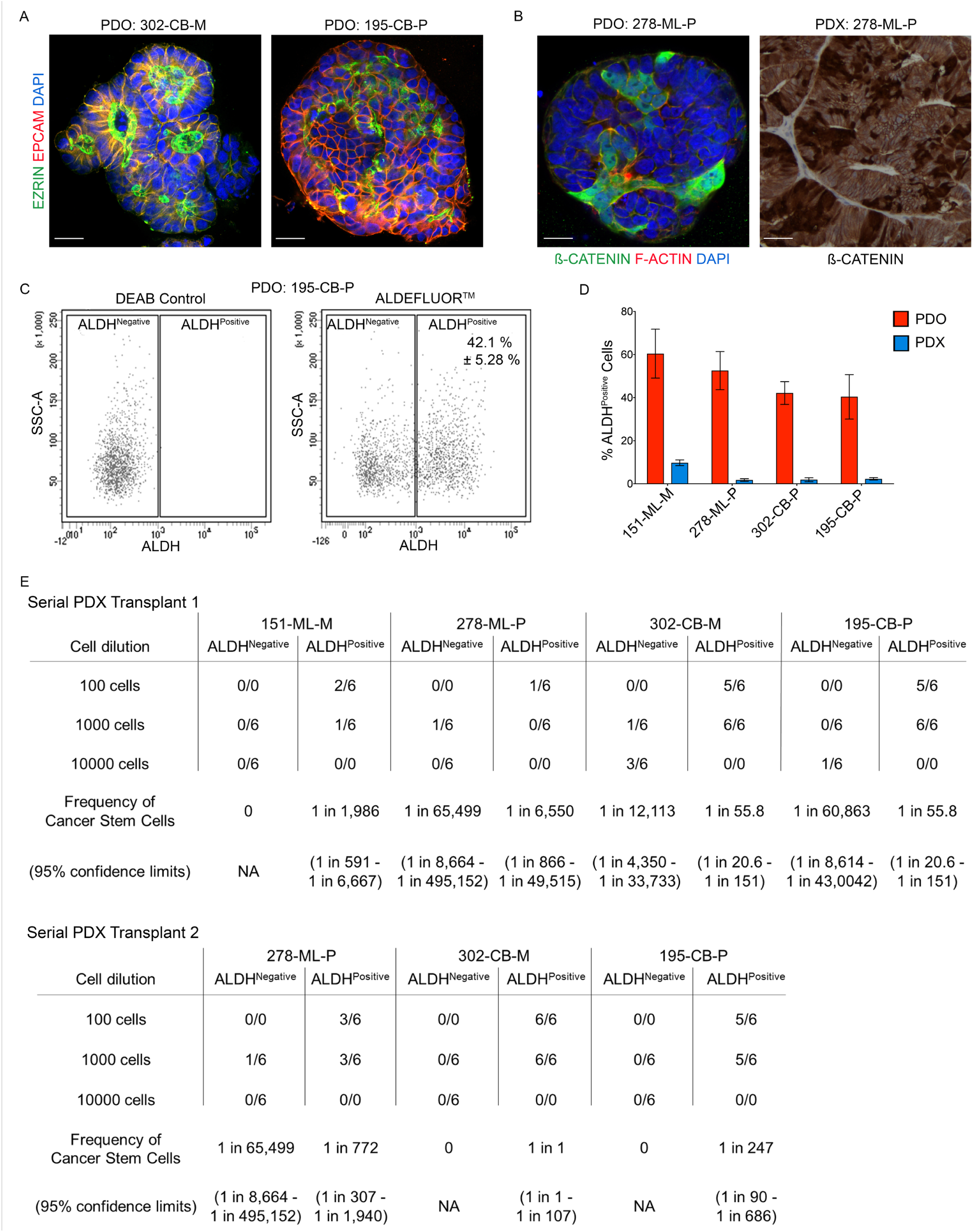
Colon cancer PDOs are heterogeneous and enriched for self-renewing ALDH^Positive^ CSCs. (A) Immunofluorescence staining of colon cancer PDOs for EZRIN (green) and EPCAM (red). Nuclei are stained blue with DAPI (Bars = 20 µm). (B) Immunofluorescence staining of a PDO for BETA-CATENIN (green) and F-ACTIN (red) (left hand side) and immunostaining of a PDX model for BETA-CATENIN (right hand side) (Bars = 20 µm). (C) Representative Aldefluor Assay FACS plots of cells derived from PDO model 195-CB-P (data from 10 independent experiments). DEAB (diethylaminobenzaldehyde) is a specific inhibitor of ALDH and is used to control for background fluorescence. (D) Frequency (±SD) of ALDH^Positive^ cells in PDOs and corresponding PDX models (data from 10 independent experiments). (E) Tables show results of two rounds of limiting dilution serial xenotransplantation of ALDH^Positive^ and ALDH^Negative^ cells from previously established PDO derived xenograft models. The number of successfully established tumors as a fraction of the number of animals transplanted is given. P-values for pairwise tests of differences in CSC frequencies between ALDH^Positive^ versus ALDH^Negative^ cells in 151-ML-M, 278-ML-P, 302-CB-M and 195-CB-P in serial transplant round one tumors are 1.12 × 10^−4^, 1.37 × 10^−1^, 8.39 × 10^−14^ and 2.92 × 10^−17^ respectively and in 278-ML-P, 302-CB-M and 195-CB-P serial transplant round two tumors are 3.82 × 10^−7^. 3.67 × 10^−22^ and 3.78 × 10^−15^, respectively. (See also Figure S1 and Table S1).

### Colon CSCs are enriched for embryonic and nervous system development gene expression signatures

In order to identify modulators of colon CSCs, ALDH^Negative^ cells and ALDH^Positive^ CSCs were isolated from PDO and PDX models and subjected to whole transcriptome analysis by RNA-sequencing. *ALDH1A1* is a marker of poor prognosis in several cancer types^27,29,45–49^ and has been reported to be responsible for the aldehyde dehydrogenase activity that defines the ALDH^Positive^ cell fraction in the Aldefluor™ assay^50^. However, nineteen different isoforms of ALDH exist and several of these, including *ALDH1A2, ALDH1A3* and *ALDH2* have also been reported to be involved in the Aldefluor™ assay^51– 53^. Here we show that *ALDH1A1* expression is enriched in ALDH^Positive^ CSCs compared to ALDH^Negative^ cells (Figure 2A and S1). GSEA of ALDH^Positive^ and ALDH^Negative^ cells isolated from PDO and PDX models demonstrated that ALDH^Positive^ CSCs are enriched for nervous system development, TNF*α* via NF*κ*B signaling, epithelial mesenchymal transition (EMT), embryonic development and Wnt signaling transcripts (Figure 2B).

**Figure 2.**
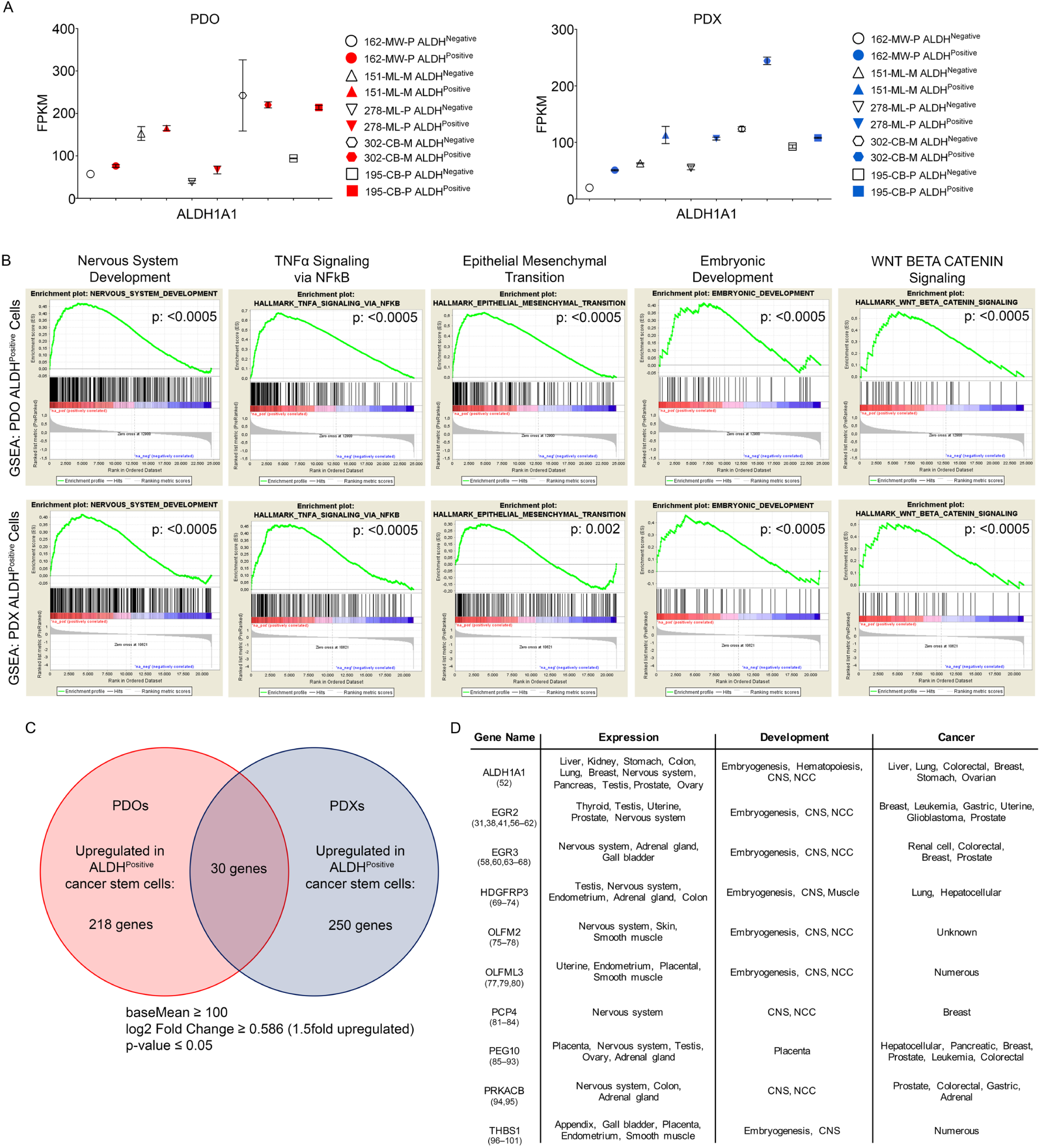
PDO and PDX ALDH^Positive^ CSCs are enriched for nervous system development gene sets and neural crest stem cell genes. (A) RNA sequencing generated FPKM values for ALDH1A1 (n = 3 separate cell preparations). (B) Gene set enrichment analysis for nervous system development (nominal p-values = <0.0005), TNFα signaling via NFkB (nominal p-value = <0.0005), epithelial to mesenchymal transition (nominal p-values = <0.0005 and 0.002), embryonic development (nominal p-value = <0.0005), and Wnt β-Catenin signaling (nominal p-values= <0.0005) in ALDH^Positive^ cells (compared to ALDH^Negative^ cells) from PDO models (top panels) and PDX models (bottom panels). (C) Venn diagram shows the number of RNA-sequencing generated transcripts upregulated in PDO ALDH^Positive^ cells (218 genes) and PDX ALDH^Positive^ cells (250 genes) and upregulated in both PDO ALDH^Positive^ cells and PDX ALDH^Positive^ cells (30 genes) n = 4 separate cell preparations, basemean greater than or equal to 100, log2 fold change = 1.5 fold upregulated, p-value <0.05). (D) Table shows 10 genes upregulated in both PDO ALDH^Positive^ cells and PDX ALDH^Positive^ cells selected for functional analysis by RNA-interference (relevant literature is cited in brackets below gene names). (See also Figure S2 and S3).

Differential gene expression analysis identified 218 genes upregulated in PDOs and 250 genes upregulated in PDX models compared to ALDH^Negative^ cells. Of these, 30 genes were found to be differentially expressed in both ALDH^Positive^ PDO and PDX cells (Figure 2C). Interestingly, many of these differentially expressed and common PDO-PDX genes are expressed during embryogenesis and have a role in neural crest cell (NCC) and central nervous system (CNS) development. Of these 30 common genes (Figure S2) 10, *ALDH1A1*^50^, *EGR2*^31,38,41,54–60^, *EGR3*^56,58,61–66^, *HDGFRP3*^67–72^, *OLFM2*^73–76^, *OLFML3*^75,77,78^, *PCP4*^79–82^, *PEG10*^83–91^, *PRKACB*^92,93^, and *THBS1*^94–99^, were selected for functional analysis based on their tissue expression and roles in development and cancer (Figure 2D, S2 and S3).

### *EGR2* is required for colon CSC survival in non-adherent cell culture

The ability of CSCs to survive and form spheroids in non-adherent cell culture is the gold standard assay for the assessment of normal stem cells and CSCs *in vitro*^100,101^. In order to test 10 of the differentially expressed genes common to ALDH^Positive^ PDO-PDX models, cells were transfected with siRNAs against *ALDH1A1, EGR2, EGR3, HDGFRP3, OLFM2, OLFML3, PCP4, PEG10, PRKACB* and *THBS1* (Figure 3B), serially plated at limiting dilution into low-attachment plates and assessed for spheroid formation. siRNA *EGR2* caused a significant decrease in spheroid formation and proliferation in all models (Figure 3A, C and D). Immunostaining of PDO, PDX and clinical samples demonstrated *EGR2* to be ubiquitously expressed, with increased cytoplasmic and nuclear expression in cancer compared to normal mucosa (Figure S4).

**Figure 3.**
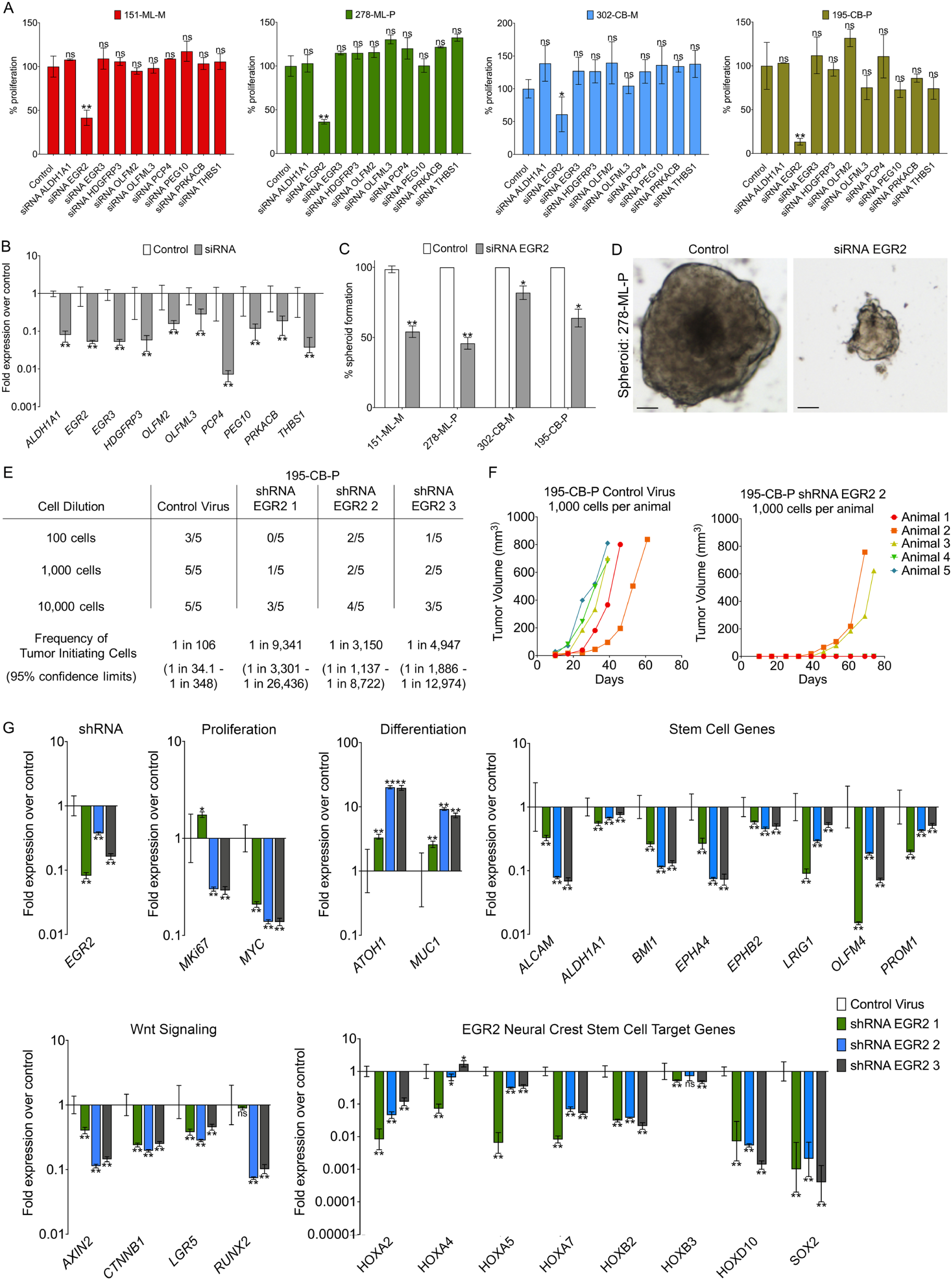
*EGR2* is required for CSC tumorigenicity and differentiation and regulates expression of NCSC *HOX* genes and *SOX2*. (A) Proliferation of siRNA transfected patient-derived colon cancer cells in non-adherent cell culture compared to control cells (mean ± SD; data from three independent experiments). *p-value < 0.05; ***p-value < 0.001 (t test). (B) Fold expression of *ALDH1A1, EGR2, EGR3, HDGFRP3 OLFML2, PCP4, PEG10, PRKACB* and *THBS1* RT-PCR gene expression data (±95% confidence intervals) in siRNA transfected 278-ML-P cells (n=3 independent cell preparations) over the comparator population (control siRNA transfected 278-ML-P cells) (see also Table S2 and S4). (C) Frequency of siRNA EGR2 spheroid formation in non-adherent cell culture compared to control transfected cells (mean ± SD; data from three independent experiments). ns = not significant; *p-value < 0.05; **p-value < 0.01 (t test). (D) Representative images of a 278-ML-P control spheroid (LHS) and a siRNA EGR2 spheroid (RHS) in non-adherent cell culture (Bars = 100 µm). (E) Table shows results of limiting dilution transplantation of control virus transduced and shRNA EGR2 transduced 195-CB-P cells. The number of established tumors as a fraction of the number of animals transplanted is given. P-values for pairwise tests of differences in CSC frequencies between control virus versus shRNA EGR2 1, shRNA EGR2 2 and shRNA EGR2 3 195-CB-P cells are 6.9 ×10^−9^, 4.9 × 10^−6^ and 6.92 × 10^−8^, respectively. (F) Growth curves for xenografts derived from control virus transduced cells and shRNA EGR2 transduced cells. (G) Fold expression of *EGR2*, proliferation, differentiation, stem cell genes, Wnt signaling and EGR2 NCSC target genes RT-PCR gene expression data (±95% confidence intervals) in four separate 195-CB-P shRNA EGR2 tumors over the comparator population (four control virus transduced 195-CB-P xenografts). Significant differences are as follows: ∗p < 0.05, ∗∗p < 0.01. (see also Table S3).

### shRNA EGR2 cells are less tumorigenic, more differentiated and have decreased expression of *HOX* genes and *SOX2*

Limiting dilution xenotransplantation of control virus transduced and shRNA EGR2 transduced 195-CB-P cells was carried out to determine if *EGR2* regulates tumorigenesis *in vivo*. Control virus transduced cells generated xenografts at each cell dilution tested but shRNA EGR2 transduced cells were significantly impaired in their ability to generate tumors when transplanted at low cell number (Figure 3E). In addition, shRNA EGR2 tumors grew more slowly than control transduced cells (Figure 3F). These data demonstrate that loss of *EGR2* in CSCs significantly decreased their tumorigenic capacity. Quantitative RT-PCR analysis of three shRNA EGR2 tumors confirmed that the shRNA EGR2 knockdown was present (Figure 3G). Significantly, expression of proliferation (*MKI67, MYC*), intestinal stem cell genes (*ALCAM, ALDH1A1, BMI1, EPHA4, EPHB2, LRIG1, OLFM4, PROM1*) and Wnt signaling genes (*AXIN2, CTNNB1, LGR5, RUNX2*) were decreased, while the expression of differentiation markers, including the tumor suppressor and Wnt signaling target *ATOH1*, were strongly increased (Figure 3G). Interestingly, *ATOH1* is also essential for neuronal differentiation during embryonic development^102–109^.

During embryogenesis *EGR2* has a conserved role in regulating embryonic master transcriptional regulator HOX genes and the stem cell regulator *SOX2*^30–32,34–41^. In addition, several HOX genes and *SOX2* have recently been shown to be enriched in and to regulate colon CSCs^17–20,23^. We therefore investigated whether these genes were similarly regulated by *EGR2* in colon PDX tumors. Notably, we found that *SOX2* and several *HOX* genes, namely *HOXA2, HOXA4, HOXA5, HOXA7, HOXB2, HOXB3* and *HOXD10*, were downregulated in shRNA EGR2 tumors (Figure 3G).

### *EGR2, ATOH1, HOXA2, HOXA4, HOXA5, HOXA7, HOXB2* and *HOXB3* are predictors of patient outcome in colorectal cancer

To characterize *EGR2, ATOH1, HOXA2, HOXA4, HOXA5, HOXA7, HOXB2, HOXB3 HOXD10* and *SOX2* expression in clinical samples, we analyzed expression across different colorectal tumor stages (Figure 4A). These data demonstrated that *EGR2* (p-value 0.027), *HOXA2* (p-value 0.026), *HOXA4* (p-value 0.000075) *HOXA5* (p-value 0.001), *HOXA7* (p-value 0.009), *HOXB3* (p-value 0.0016) and HOXD10 (p-value 0.043) expression are more enhanced in late stage T4 clinical tumors. Of these, *HOXA4, HOXA5, HOXA7*, and *HOXB3* are significant at FDR < 5%. Analysis of Kaplan-Meier survival curves showed that patients with higher *EGR2, HOXA2, HOXA4, HOXA5, HOXA7, HOXB2 and HOXB3* expression have a poorer clinical outcome (p-values 0.00017, 0.0028, 0.0006, 0.0043, 0.0022, 0.00025 and 0.019, respectively). Of these, higher *EGR2, HOXA2, HOXA4, HOXA5* and *HOXA7* are significant at FDR < 5%. Furthermore, these data demonstrated that high levels of *ATOH1* are predictive of good prognosis (p-value 0.0013). These data support *ATOH1, EGR2* and its target genes *HOXA2, HOXA4, HOXA5, HOXA7 and HOXB3* as potential new biomarkers for CRC prognosis.

**Figure 4.**
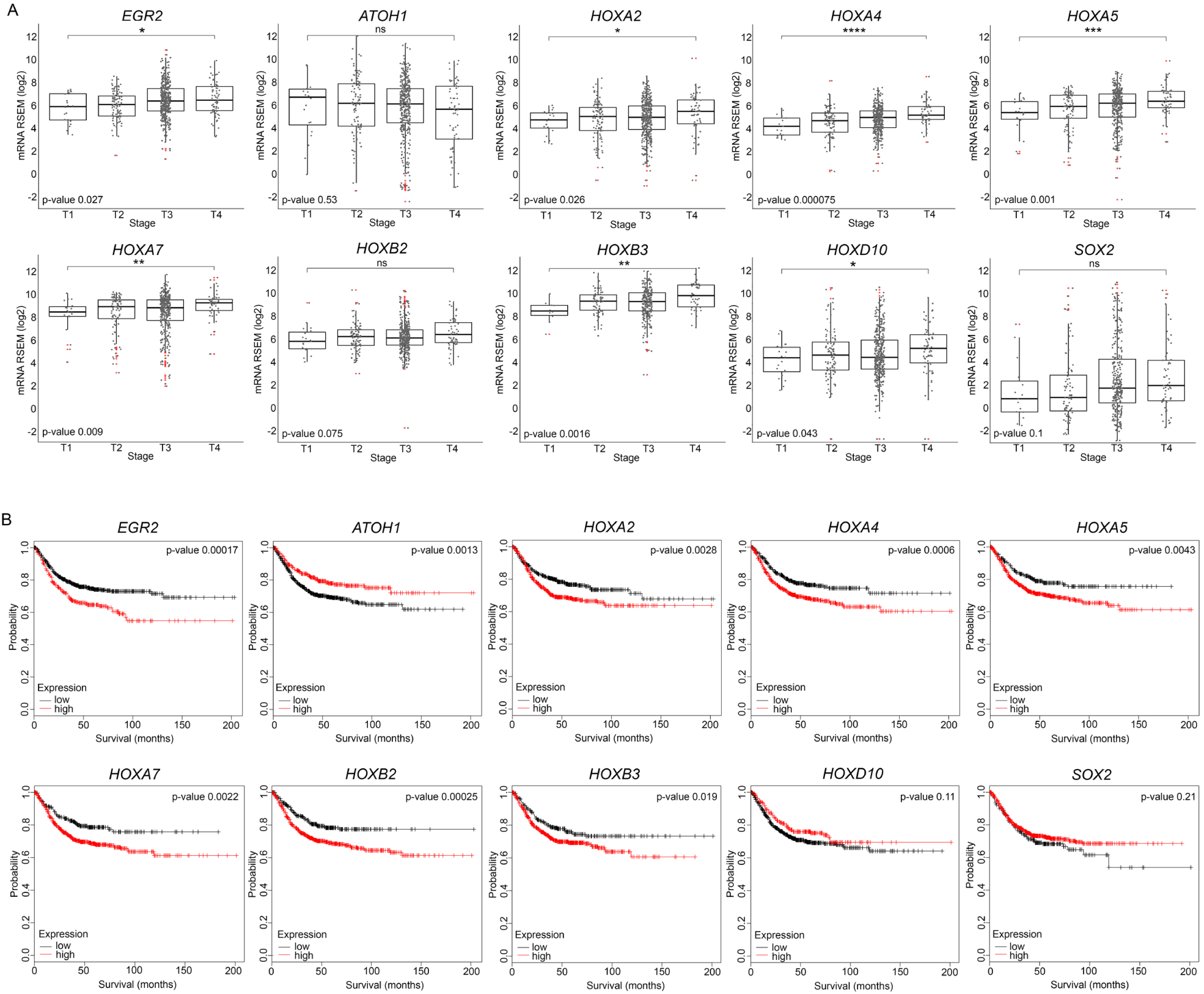
*EGR2, HOXA2, HOXA4, HOXA5, HOXA7* and *HOXB3* are increased in late stage tumors and are indicators of poor prognosis in clinical samples. (A) Expression *of EGR2, ATOH1, HOXA2, HOXA4, HOXA5, HOXA7, HOXB2, HOXB3, HOXD10* and *SOX2* in colorectal cancer patients across different tumor stages (T1 v T4, p-value = 0.027, 0.53, 0.026, 0.000075, 0.001, 0.009, 0.075, 0.0016, 0.043 and 0.1, respectively). Of these, *HOXA4, HOXA5, HOXA7*, and *HOXB3* are significant at FDR < 5%. RNAseq and clinical data of 533 patients (n=378 colon adenocarcinoma, n=155 rectal adenocarcinoma) was extracted from cBioPortal. (B) Kaplan-Meier survival curves for *EGR2, ATOH1, HOXA2, HOXA4, HOXA5, HOXA7, HOXB2, HOXB3, HOXD10* and *SOX2* in colorectal cancer patients comparing lower third percentile to upper third percentile (logrank p-values = 0.00017, 0.0013, 0.0028, 0.0006, 0.0043, 0.0022, 0.00025, 0.019, 0.11 and 0.21, respectively. Of these, higher *EGR2, HOXA2, HOXA4, HOXA5* and *HOXA7* are significant at FDR < 5%. Results based upon data generated by the Kaplan-Meier Plotter (www.kmplot.com/analysis)^194^.

## DISCUSSION

We previously demonstrated that colon cancer PDOs are enriched for CSCs and preserve the functional and molecular heterogeneity found *in vivo*, thus making them excellent models for the study of CSCs^43^. However, the defined conditions of the PDO culture media results in reduced cell type diversity^42^. Conversely, the *in vivo* environment promotes differentiation and reduces CSCs to a minority population. Therefore, in order to identify genes that regulate CSC survival and differentiation we carried out whole transcriptome analyses of functionally defined ALDH^Negative^ cells and ALDH^Positive^ CSCs from colon cancer PDO and PDX models and performed functional analyses of genes differentially expressed and common to ALDH^Positive^ CSCs from both models.

Interestingly, these analyses revealed transcripts associated with nervous system development and NCSCs to be highly enriched in both PDO and PDX CSCs. Recent studies have demonstrated that solid tumors, including CRC, contain nerve fibers that promote tumor growth and metastasis, indeed, neurogenesis in CRC is an independent indicator of poor clinical outcome^110,111^, but their origin and mechanism of innervation is unknown^112–117^.

A growing body of evidence has demonstrated a gut-neural axis^118–123^ in which various intestinal cells, including stem cells, interact with the autonomic nervous system (ANS), either directly^124–130^ or via the enteric nervous system (ENS)^131–133^, a network of neurons and glia within the bowel wall that regulates most aspects of intestinal function^134^, to control stem cell proliferation and differentiation^135,136^. For example, ISCs express ANS-associated alpha2A adrenoreceptor (Adra2a) and acetylcholine (ACh) receptors implicated in controlling intestinal epithelial proliferation^130,137–140^. In addition, differentiated cell types, such as intestinal enterochromaffin (EC) cells have been found to be electrically excitable and modulate serotonin-sensitive primary afferent nerve fibers via synaptic connections, enabling them to detect and transduce environmental, metabolic, and homeostatic information from the gut directly to the nervous system^141^. Recent studies have also demonstrated that enteroendocrine cells form neuroepithelial circuits by directly synapsing with vagal neurons and called for a renaming of these cells from enteroendocrine to neuropod cells^129,142^. Neuropod cells and EC cells, like all differentiated intestinal cells (enteroendocrine, enterocyte, goblet, paneth) and CSCs, derive from multipotent Lgr5^Positive^ crypt stem cells^143,144^. Significantly, colorectal CSCs themselves have been shown to be capable of generating neurons when transplanted intraperitoneally in nude mice^145^. Intestinal stem cells and CSCs should therefore possess the capacity to express nervous system genes, since they are the progenitors of cells with neural function. However, until now, no previous study had directly reported nervous system gene enrichment in ISCs or CSCs.

We therefore carried out gene ontology analysis of Lgr5^Positive^ crypt stem cell transcriptomes from earlier studies^146–148^. In agreement with our CSC data (Figure 2), this analysis revealed normal ISCs to also be enriched for nervous system genes (Figure S5). In addition, the PDOs showed ubiquitous staining for the epithelial cell marker EPCAM (Figure 1A), demonstrating that they do not contain a separate non-epithelial neural cell lineage that could be the origin of the nervous system gene expression. Overall, these data suggest that CSCs may be a source of the neural connections that interact with the ANS and peripheral nervous system (PNS) to drive tumor growth and metastasis^112–116^. Denervation of the ANS and PNS, which causes loss of autonomic neurotransmitters in the gut, results in loss of crypt stem cell proliferation and suppression of tumorigenesis^124– 128,131,132,149–151^. The inhibition of nervous system gene transcription in CSCs and their progeny may therefore provide a novel therapeutic strategy in colorectal cancer, with results similar to denervation^149,150,152^.

During embryonic development, the PNS, of which the ENS is a part, arises from NCSCs, multipotent and highly migratory stem cells that move throughout the embryo to colonize multiple organ primordia and differentiate into numerous cell types^24,153–155^. Recently, self-renewing NCSCs have been discovered in post-natal tissue^156–160^, including the adult gut^161,162^, although the degree to which these cells contribute to the adult tissue is not yet known.

*EGR2* is a conserved regulator and marker of NCSCs that acts upstream of several *HOX* genes and *SOX2* to control cell fate in embryonic and nervous system stem cells^30–41^. Interestingly, its expression is also rapidly activated after wounding in the embryonic and adult mouse^33^, suggesting a role in adult tissue stem cells, which contribute to tissue regeneration and wound repair^163^. However, no previous study has identified a role for *EGR2* in CRC. Here, we demonstrate that *EGR2* is enriched in colon CSCs and is required for tumorigenicity and to maintain CSCs in an undifferentiated state by regulating *HOX* genes and *SOX2*.

*SOX2* is one of the early genes activated in the developing neural crest and has a broad role as a transcriptional regulator in embryonic and adult stem cells^15,164–169^. In embryonic and adult neural stem cells, it is required for the maintenance of neural stem cell properties, including proliferation, survival, self-renewal and neurogenesis^170–174^. In the intestine, its expression results in cell fate conversion and redirects the intestinal epithelium to a more undifferentiated phenotype^175–177^. In addition, *SOX2* has been associated with a stem cell state in several cancer types^178–180^ and is aberrantly expressed in CRC^176,181,182^. Overall, these data, combined with our own, support a role for *SOX2* in CRC tumor initiation and progression, possible by promoting neural specification in CSCs and their descendants.

*HOX* genes have been reported to be enriched in and required for the maintenance of normal stem cells and CSCs in various adult tissues^11,13,16,183–189^. Recently, *HOXA4, HOXA9* and *HOXD10* were shown to be selectively expressed in ALDH^Positive^ intestinal crypt stem cells and colon CSCs, to promote self-renewal and regulate expression of stem cell markers^17,18^. Here, we demonstrate that the same *HOX* genes that are regulated by *EGR2* in NCSCs are also regulated by *EGR2* in colon CSCs and that several of these, *HOXA2, HOXA4, HOXA5, HOXA7, HOXB2, HOXB3*, along with *EGR2*, are indicators of poor prognosis in CRC.

These data demonstrate that colon CSCs are enriched for neural crest and nervous system development genes, including the NCSC regulator *EGR2*, which controls *SOX2* and *HOX* genes to maintain CSCs in an undifferentiated state required for tumorigenesis. Targeting *EGR2* to induce differentiation and block potential intestinal-neural cell specification, e.g. by downregulating the neural stem cell regulator SOX2, may offer a novel therapeutic strategy to eliminate colon CSCs and prevent nervous system driven proliferation and metastasis.

## EXPERIMENTAL PROCEDURES

### Human tissue samples and establishment of patient-derived cancer organoid cell cultures

Tumor material was obtained with informed consent from CRC patients under approval from the local Institutional Review Board of Charité University Medicine (Charité Ethics Cie: Charitéplatz 1, 10117 Berlin, Germany) (EA 1/069/11) and the ethics committee of the Medical University of Graz and the ethics committee of the St John of God Hospital Graz (23-015 ex 10/11). Tumor staging was carried out by experienced and board-certified pathologists (Table S1). Cancer organoid cultures were established and propagated as described^42,44^.

### Limiting dilution xenotransplantation

Housing and handling of animals followed European and German Guidelines for Laboratory Animal Welfare. Animal experiments were conducted in accordance with animal welfare law, approved by local authorities, and in accordance with the ethical guidelines of Bayer AG. PDO derived PDX models were processed to single cells and sorted by FACS (BD FACS Aria II) for ALDH activity (Aldefluor assay) and DAPI to exclude dead cells. Cells were then re-transplanted at limiting dilutions by injected subcutaneously in PBS and Matrigel (1:1 ratio) at limiting cell dilutions into female 8 – 10-week-old nude^-/-^ mice.

### Histology and immunohistochemistry

Tumors were fixed in 4% paraformaldehyde overnight for routine histological analysis and immunohistochemistry. Immunohistochemistry was carried out via standard techniques with non-phospho (Active) β-Catenin (#8814, rabbit monoclonal, Cell Signaling Technology; diluted 1:200) and EGR2 (ab43020, Abcam, rabbit IgG, polyclonal, diluted 1:1000) antibodies. Negative controls were performed using the same protocols with substitution of the primary antibody with IgG-matched controls (ab172730, rabbit IgG, monoclonal [EPR25A], Abcam). Colorectal cancer tissue microarrays from the OncoTrack patient cohort^44^ were obtained from The Institute of Pathology, Medical University Graz, Austria and analyzed using Aperio TMALab and Image software (Leica Biosystems).

### Immunofluorescence staining of PDOs

For immunofluorescence imaging, cancer organoid cultures were fixed in 4% paraformaldehyde for 30 min at room temperature and permeabilized with 0.1% Triton X-100 for 30 min and blocked in phosphate-buffered saline (PBS) with 10% bovine serum albumin (BSA). Samples were incubated with primary antibodies overnight at 4°C. Antibodies used were Non-phospho (Active) β-Catenin (#8814, rabbit monoclonal, Cell Signaling Technology; diluted 1:200), EZRIN (ab40839, rabbit monoclonal, Abcam, diluted 1:200), EPCAM (#2929, mouse monoclonal, Cell Signaling Technology, diluted 1:500) and EGR2 (ab43020, rabbit polyclonal, Abcam, diluted 1:1000). Samples were stained with a conjugated secondary antibody overnight at 4°C. F-actin was stained with Alexa Fluor® 647 Phalloidin (#A22287, Thermo Fisher; diluted 1:20) for 30 min at room temperature. Nuclei were counterstained with DAPI. Negative controls were performed using the same protocol with substitution of the primary antibody with IgG-matched controls. Cancer organoids were then transferred to microscope slides for examination using a Zeiss LSM 700 Laser Scanning Microscope.

### Aldefluor Assay

Organoids and xenografts were processed to single cells and labelled using the Aldefluor Assay according to manufacturer’s (Stemcell Technologies) instructions. ALDH levels were assessed by FACS on a BD LSR II analyzer.

### RNA Sequencing

Cells were lysed in RLT buffer and processed for RNA using the RNeasy Mini Plus RNA extraction kit (Qiagen). Samples were processed using Illumina’s TrueSeq RNA protocol and sequenced on an Illumina HiSeq 2500 machine as 2×125nt paired-end reads. The raw data in Fastq format were checked for sample quality using our internal NGS QC pipeline. Reads were mapped to the human reference genome (assembly hg19) using the STAR aligner (version 2.4.2a). Total read counts per gene were computed using the program “featureCounts” (version 1.4.6-p2) in the “subread” package, with the gene annotation taken from Gencode (version 19). The “DESeq2” Bioconductor package was used for the differential-expression analysis.

### siRNA transfection

Cells were seeded in 100 µl volumes of Accell™ Delivery Media (Dharmacon™) at 1.0 × 10^5^ cells per well in ultra-low attachment 96-well plates and transfected with 2 µM concentrations of Accell™ siRNAs (Table S2) and control siRNA (Accell™ non-targeting siRNA control) (Dharmacon™) by incubating for up to 72 h in Accell siRNA Delivery Media.

### Viral transduction

Cells were seeded in 100 µl volumes of antibiotic free culture media at 1.0 ×10^5^ cells per well in ultra-low attachment 96-well plates. Control and shRNA lentiviruses were purchased from Sigma-Aldrich (Table S3). Viral particles were added at a multiplicity of infection of 1. Cells were transduced for up to 96 h or until GFP positive cells were observed before being embedded in Matrigel for the establishment of lentiviral transduced cancer organoid cultures. Puromycin (2 µg/ml) was used to keep the cells under selection.

### Limiting dilution spheroid assays

For siRNA spheroid assays, transfected live (DAPI^Negative^) cells were sorted at 10 cells per well into 96-well ultra-low attachment plates. 20 days later wells containing spheroids were counted and used to calculate CSC frequency using ELDA software. Proliferation was measured using the CellTiter-Glo^®^ Luminescent Cell Viability Assay.

### Gene expression analysis

For quantitative real-time RT-PCR analysis RNA was isolated using the RNeasy Mini Plus RNA extraction kit (Qiagen). cDNA synthesis was carried out using a Sensiscript RT kit (Qiagen). RNA was transcribed into cDNA using an oligo dTn primer (Promega) per reaction. Gene expression analysis was performed using TaqMan® Gene Expression Assays (Applied Biosystems) (Table S4) on an ABI Prism 7900HT sequence detection system (Applied Biosystems). GAPDH was used as an endogenous control and results were calculated using the Δ-ΔCt method. Data were expressed as the mean fold gene expression difference in three independently isolated cell preparations over a comparator sample with 95% confidence intervals. Pairwise comparison of gene expression was performed using R^190^ together with package ggplot2^191^ on log2 transformed RNAseq data from 533 patients with clinical data (n=378 colon adenocarcinomas, n=155 rectal carcinomas staged T1-T4) extracted from the cBioPortal for Cancer Genomics (cbioportal.org)^192,193^. Survival curves were generated using the Kaplan-Meier Plotter (www.kmplot.com/analysis)^194^. Gene ontology enrichment analysis was carried out using the Gene Ontology Resource (www.geneontology.org)^195,196^.

### Statistical analysis

GraphPad Prism 6.0 was used for data analysis and imaging. All data are presented as the means ± SD, followed by determining significant differences using the two-tailed t test. Significance of RT-PCR data was determined by inspection of error bars as described by Cumming *et al*. (2007)^197^. Limiting-dilution frequency and probability estimates were analyzed by the single-hit Poisson model and pairwise tests for differences in stem cell frequencies using the ELDA software (http://bioinf.wehi.edu.au/software/elda/index.html, Hu and Smyth, 2009)^198^. Gene set enrichment analysis was carried out using pre-ranked feature of the Broad Institute GSEA software version 2 using msigdb v5.1 gene sets^199,200^. The ranking list was derived from the fold changes (1.5fold upregulated) calculated from the differential gene expression calculation and nominal p-values. P-values <0.05 were considered as statistically significant. For the final list of significant genes, False Discovery Rate was computed using the Benjamini-Hochberg method^201^.

## Acknowledgements

We thank Dorothea Przybilla, and Cathrin Davies (Laboratory of Molecular Tumor Pathology, Charité Universitätsmedizin Berlin, Germany) for technical and cell culture assistance. The research leading to these results has received support from the Innovative Medicines Initiative Joint Undertaking under Grant Agreement 115234 (OncoTrack), the resources of which are composed of financial contribution from the European Union Seventh Framework Programme (FP7/2007-2013) and EFPIA companies in kind contribution. A.S., T.J., D.M. and. D.H. are employees of Bayer AG. R.L., J.T. and M.L. are employees of Nuvisan ICB GmbH. C.R.A.R. is a co-founder of CELLphenomics.

## Authors Contribution

Conceptualization, J.L.R.; Methodology, J.L.R. and M.L.; Investigation, J.L.R., D.S., S.S., A.S., R.L., J.T., T.J., J.H., N.G.-S., and M.L.; Writing, J.L.R.; Visualization, J.L.R; Data Curation, A.S., J.T.; Resources, J.H., U.K., C.R.A.R. and B.G.; Supervision, J.L.R., D.M., D.H., R.S., and M.L.

## Accession Numbers

Array data are available in the ArrayExpress database (www.ebi.ac.uk/arrayexpress) under accession numbers E-MTAB-5209 and E-MTAB-8927.

## Supplementary Information

**Figure S1:**
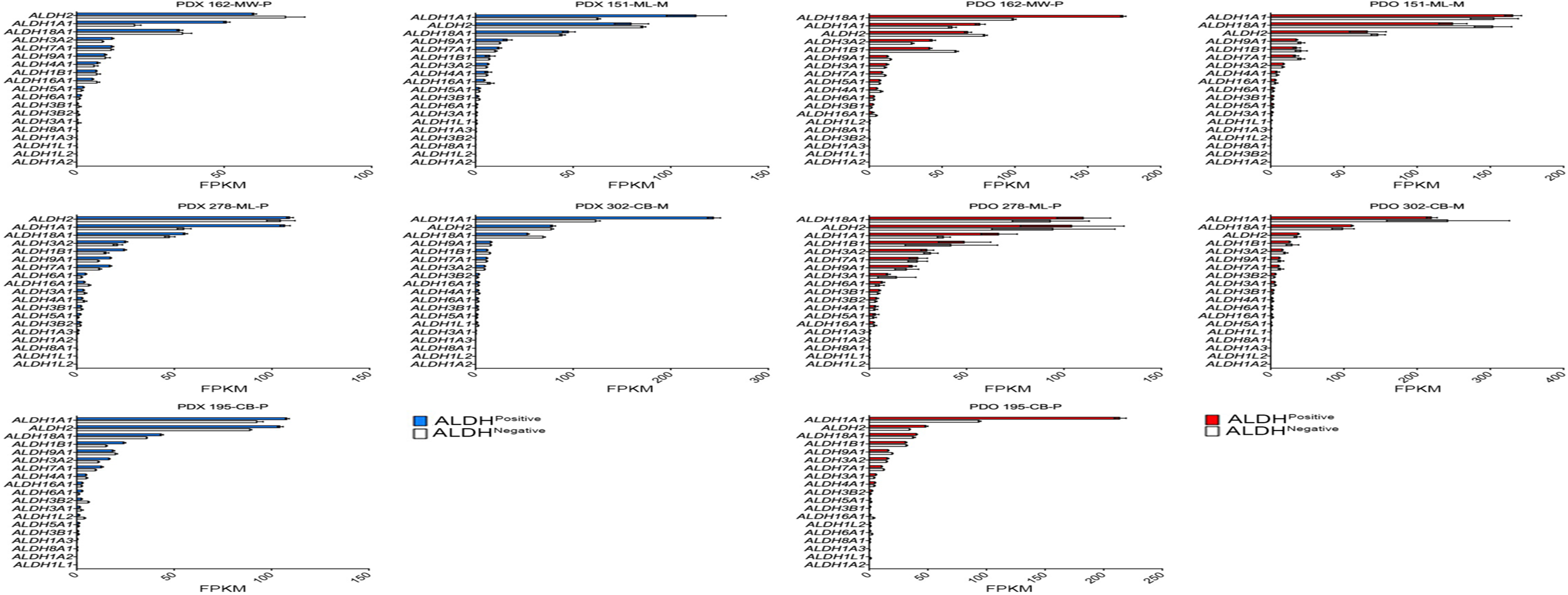
ALDH isoform expression in ALDH^Positive^ PDO (LHS) and PDX (RHS)cells. Related to Figure 1 and 2.

**Figure S2:**
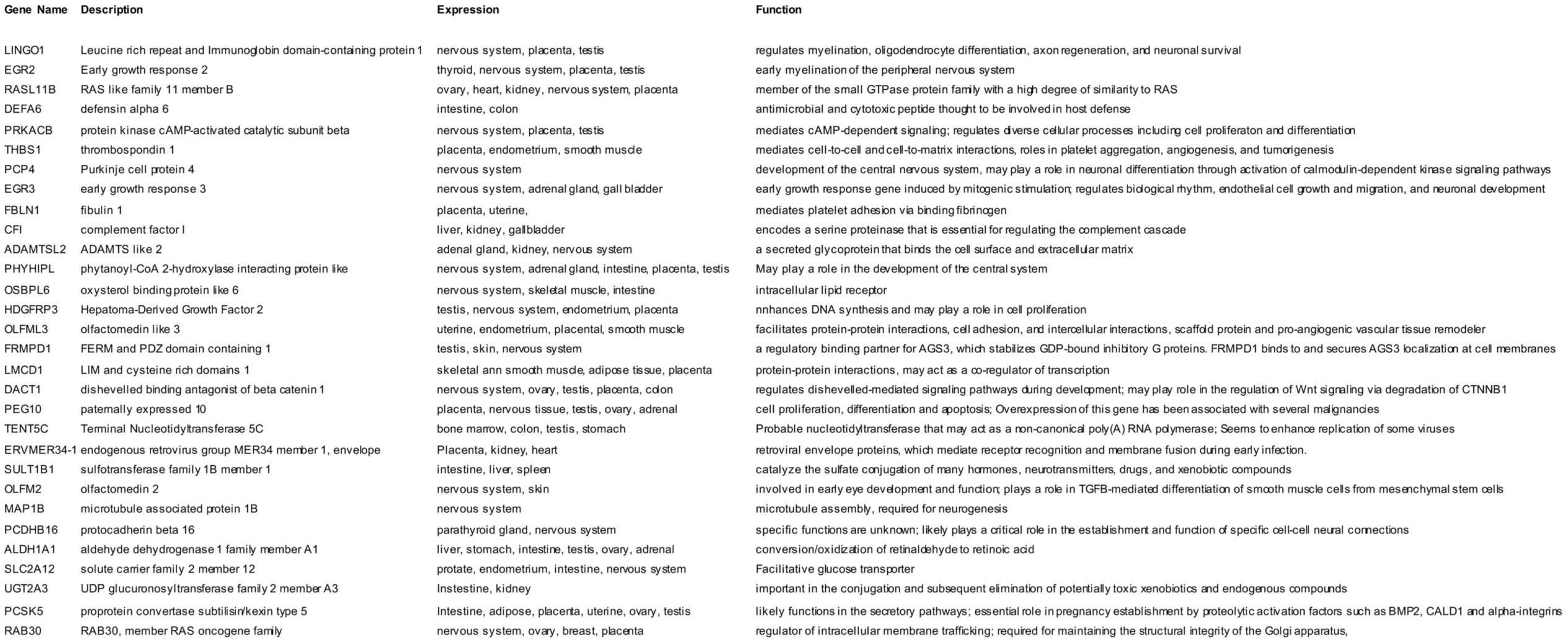
List of genes differentially expressed and common in ALDH^Positive^ PDO and ALDH^Positive^ PDX CSCs. Related to Figure 2.

**Figure S3:**
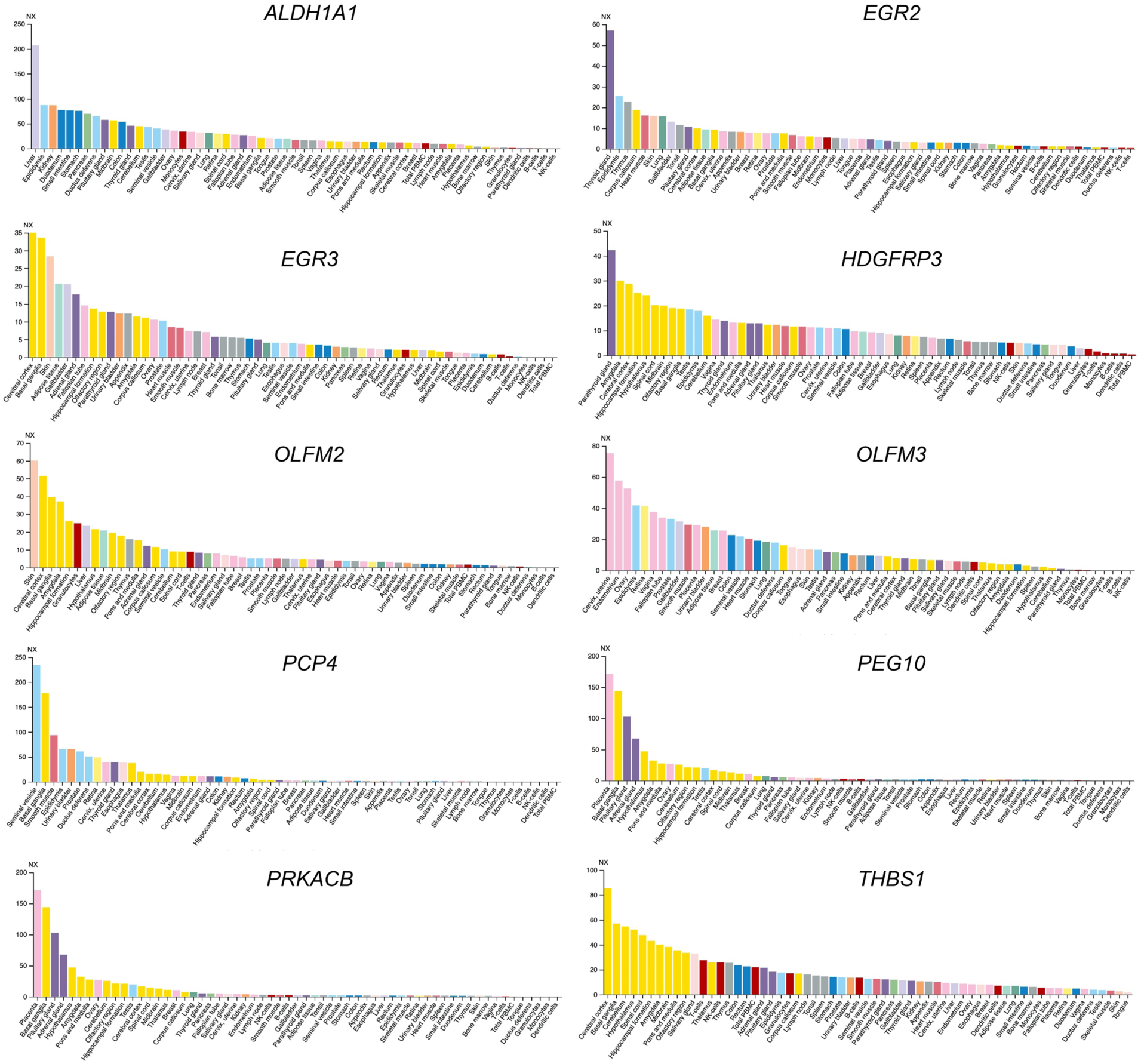
RNA tissue expression of differentially expressed and common ALDH^Positive^ PDO and ALDH^Positive^ PDX cells. Related to Figure 2. Consensus RNA-seq dataset from the Human Protein Atlas (HPA) the Genotype-Tissue Expression (GTEx) project and CAGE data from FANTOM5 project. Color-coding is based on tissue groups, each consisting of tissues with functional features in common.

**Figure S4:**
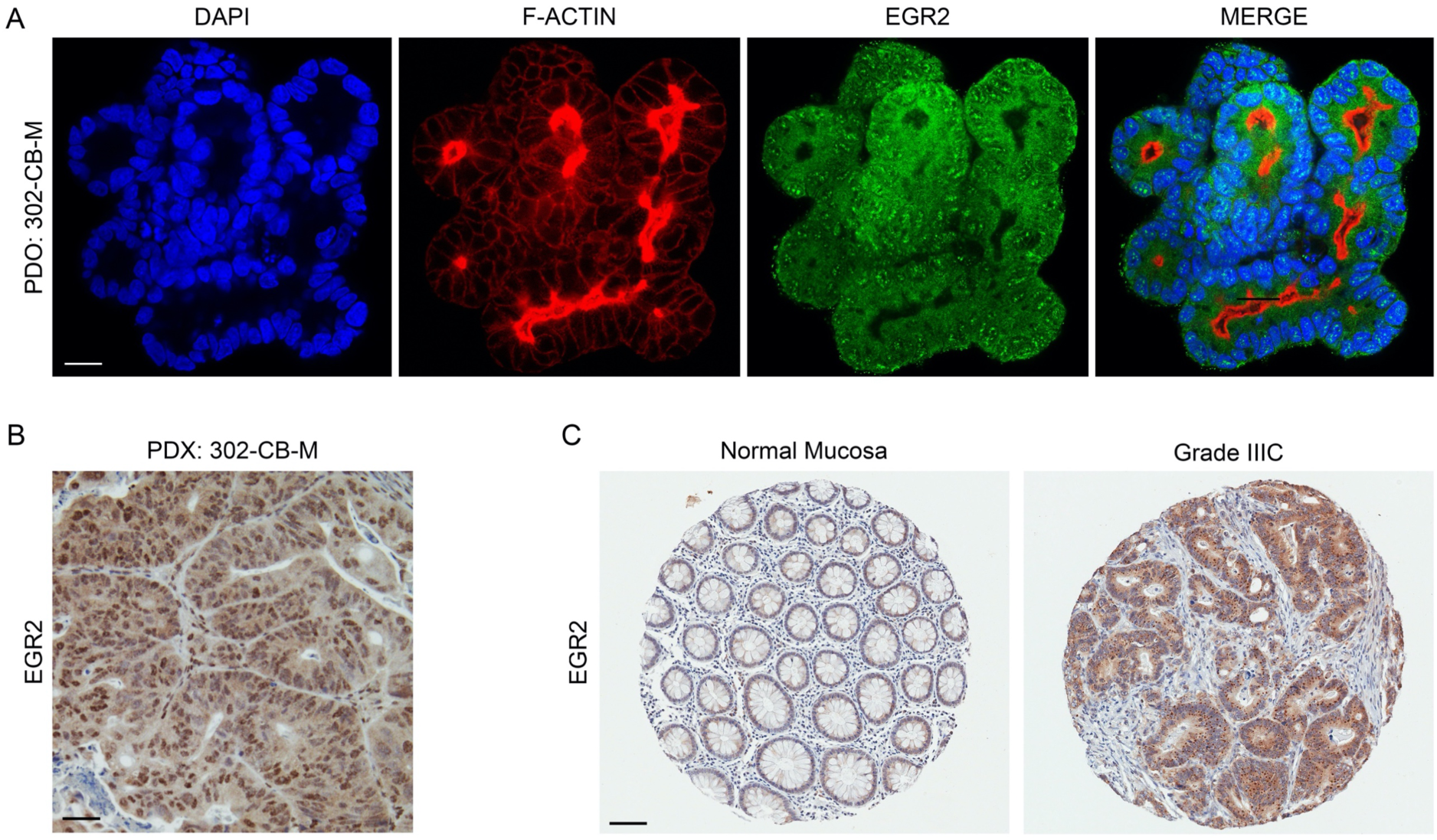
EGR2 immunostaining in PDO, PDX and clinical samples. (A) Immunofluorescence staining of PDO for EGR2 (green) and F-ACTIN (red). Nuclei are stained blue with DAPI (Bars = 20 µm). Immunostaining of PDX tissue (B) and tissue microarrays of normal intestinal mucosa and colorectal cancer patient tissue (C) for EGR2. (Bars = 200 µm). Related to Figure 3.

**Figure S5:**
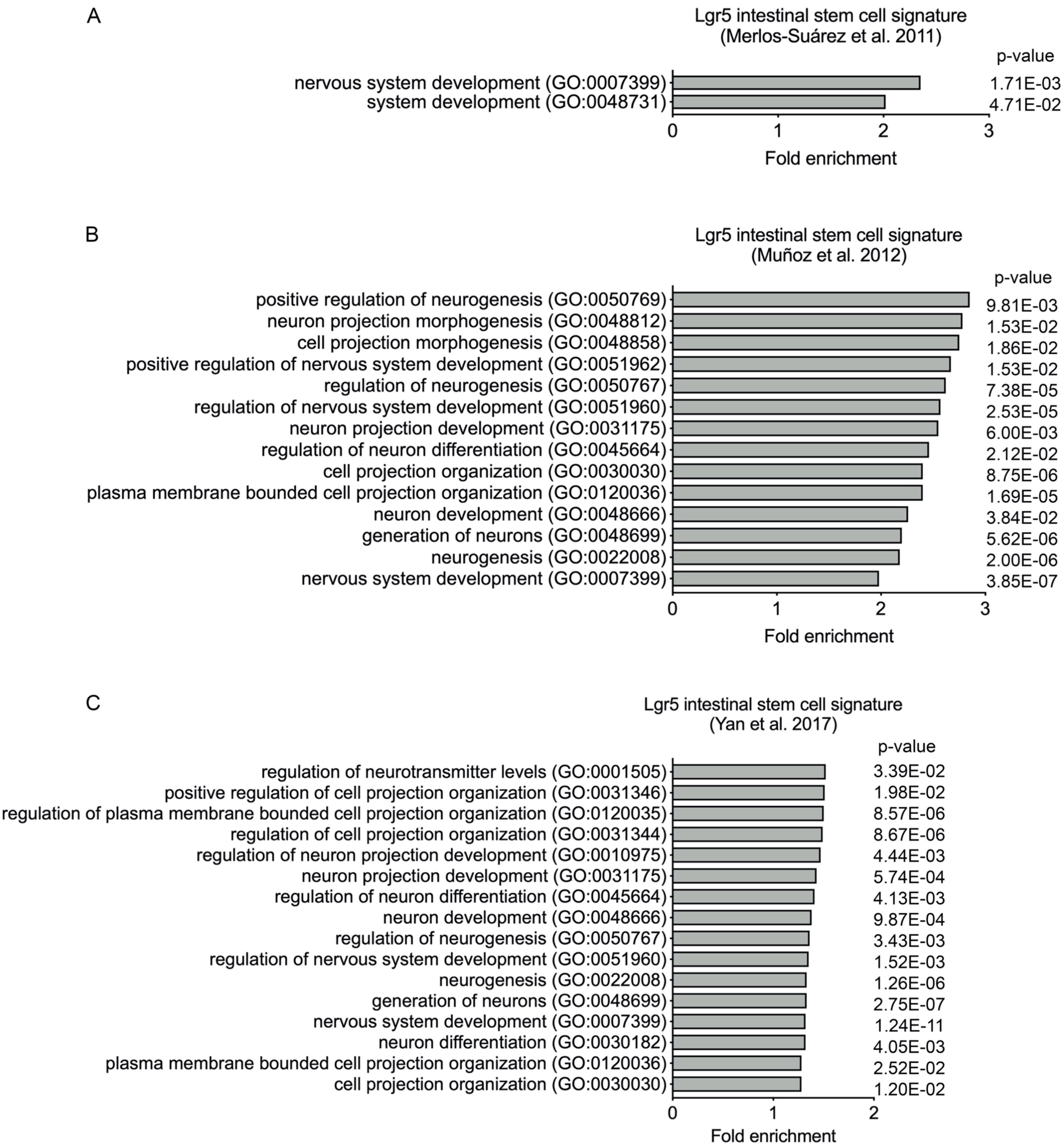
Lgr5 intestinal crypt-base stem cells are enriched for nervous system genes. Gene ontology analysis of Lgr5 intestinal stem cell gene signatures from (A) Merlos-Suarez, *et al*., 2011), (B) Munoz, *et al*., (2012) and (C) Yan, *et al*., (2017). Related to Figure 2.

**Table S1.** Tissue Origin and TNM Classification of Malignant Tumors (TNM). Related to Figure 1. T: primary tumor size, N: regional lymph nodes involved, M: distant metastasis.

| Patient Model | Origin | TNM stage | Stage |
| --- | --- | --- | --- |
| 162-MW-P | Sigmoid colon & descending colon | T3 N0 M0 | IIA |
| 151-ML-M | Liver | T2 N0 M0 , M1a | IVA |
| 278-ML-P | Sigmoid colon & descending colon | T4a N0 M0 | IIB |
| 302-CB-M | Liver | T3 N1a M1a | IVA |
| 195-CB-P | Sigmoid colon | T4a N2b M1a | IVA |

**Table S2:** Dharmacon™ Smartpool siRNAs. Related to Figure 3.

| siRNA | Dharmacon™ Product | Product Number |
| --- | --- | --- |
| ALDH1A1 | Accell Human (216) siRNA - SMARTpool | E-008722-00-5 |
| EGR2 | Accell Human (1959) siRNA - SMARTpool | E-006527-01-5 |
| EGR3 | Accell Human (1960) siRNA - SMARTpool | E-006528-00-5 |
| HDGFRP3 | Accell Human (50810) siRNA - SMARTpool | E-017093-00-5 |
| OLFM2 | Accell Human (93145) siRNA - SMARTpool | E-015212-00-5 |
| OLFML3 | Accell Human (56944) siRNA - SMARTpool | E-020325-00-5 |
| PCP4 | Accell Human (5121) siRNA - SMARTpool | E-020122-00-5 |
| PEG10 | Accell Human (23089) siRNA - SMARTpool | E-032579-00-5 |
| PRKACB | Accell Human (5567) siRNA - SMARTpool | E-004650-00-5 |
| THBS1 | Accell Human (7057) siRNA - SMARTpool | E-019743-00-5 |

**Table S3.** Lentiviral Transduction Particles. Related to Figure 3.

| LENTIVIRUS | SIGMA PRODUCT | PRODUCT NAME | VECTOR | TRC NUMBER |
| --- | --- | --- | --- | --- |
| Control | SHC003V | MISSION® tGFP™ Positive Control Transduction Particles | –pLKO.1-puro-CMV-tGFP | NA |
| shEGR2 1 | SHCLNV-NM_000399 | EGR2 MISSION shRNA Lentiviral Transduction Particles | –hPGK-Puro-CMV-tGFP | TRCN0000013839 |
| shEGR2 2 | SHCLNV-NM_000399 | EGR2 MISSION shRNA Lentiviral Transduction Particles | –hPGK-Puro-CMV-tGFP | TRCN0000013840 |
| shEGR2 3 | SHCLNV-NM_000399 | EGR2 MISSION shRNA Lentiviral Transduction Particles | –hPGK-Puro-CMV-tGFP | TRCN0000013841 |

**Table S4.** Taqman Gene Expression Assays. Related to Figure 3.

| Symbol | Gene Name | UniGene ID | TaqMan® Gene Expression Assay |
| --- | --- | --- | --- |
| ATOH1 | atonal BHLH transcription factor 1 | Hs.532680 | Hs00245453_s1 |
| AXIN2 | axin 2 | Hs.156527 | Hs00610344_m1 |
| BMI1 | BMI1 proto-oncogene, polycomb ring finger | Hs.380403 | Hs00180411_m1 |
| CTNNB1 | catenin beta 1 | Hs.476018 | Hs00355049_m1 |
| EGR2 | early growth response 2 | Hs.1395 | Hs00166165_m1 |
| EPHA4 | EPH receptor A4 | Hs.371218 | Hs00953178_m1 |
| EPHB2 | EPH receptor B2 | Hs.523329 | Hs00362096_m1 |
| GAPDH | glyceraldehyde-3-phosphate dehydrogenase | Hs.544577 | Hs02758991_g1 |
| HOXA2 | homeobox A2 | Hs.445239 | Hs00534579_m1 |
| HOXA5 | homeobox A5 | Hs.655218 | Hs00430330_m1 |
| HOXA7 | homeobox A7 | Hs.610216 | Hs00600844_m1 |
| HOXB2 | homeobox B2 | Hs.514289 | Hs01911167_s1 |
| HOXB3 | homeobox B3 | Hs.654560 | Hs05048382_s1 |
| HOXD10 | homeobox D10 | Hs.123070 | Hs00157974_m1 |
| LGR5 | leucine rich repeat containing G protein-coupled receptor 5 | Hs.658889 | Hs00969422_m1 |
| MKI67 | MKI67 | Hs.689823 | Hs04260396_g1 |
| MUC1 | mucin 1, cell surface associated | Hs.89603 | Hs00159357_m1 |
| MYC | v-myc avian myelocytomatosis viral oncogene homolog | Hs.202453 | Hs00153408_m1 |
| RUNX2 | runt related transcription factor 2 | Hs.535845 | Hs01047973_m1 |
| SOX2 | SRY-box 2 | Hs.518438 | Hs01053049_s1 |
| HDGFRP3 | hepatoma-derived growth factor, related protein 3 | Hs.513954 | Hs00274988_m1 |
| OLFM2 | olfactomedin 2 | Hs.169743 | Hs01017934_m1 |
| OLFML3 | olfactomed like 3 | Hs.9315 | Hs01113293_g1 |
| PCP4 | Purkinje cell protein 4 | Hs.80296 | Hs01113638_m1 |
| PEG10 | paternally expressed 10 | Hs.147492 | Hs00248288_s1 |
| PRKACB | protein kinase cAMP-activated catalytic subunit beta | Hs.487325 | Hs01086757_m1 |
| THBS1 | thrombospondin 1 | Hs.164226 | Hs00962908_m1 |

## Notes

https://www.ebi.ac.uk/arrayexpress/experiments/E-MTAB-8927/

https://www.ebi.ac.uk/arrayexpress/experiments/E-MTAB-5209/

### Competing Interest Statement

The authors have declared no competing interest.

